# Selective recruitment of endoplasmic reticulum-targeted and cytosolic mRNAs into membrane-associated stress granules

**DOI:** 10.1101/2021.05.12.443899

**Authors:** Jessica R. Child, Qiang Chen, David W. Reid, Sujatha Jagannathan, Christopher V. Nicchitta

**Affiliations:** Department of Cell Biology, Duke University School of Medicine, Durham, NC 27710, USA; Moderna, Inc. Cambridge, MA 02139, USA; Department of Biochemistry and Molecular Genetics, University of Colorado, Anschutz Medical Campus, Denver, CO 80045, USA; RNA Bioscience Initiative, University of Colorado, Anschutz Medical Campus, Denver, CO 80045, USA

**Keywords:** stress granule, mRNA, translational regulation, endoplasmic reticulum, unfolded protein response

## Abstract

Stress granules (SGs) are membraneless organelles composed of mRNAs and RNA binding proteins which undergo assembly in response to stress-induced inactivation of translation initiation. The biochemical criteria for mRNA recruitment into SGs are largely unknown. In general, SG recruitment is limited to a subpopulation of a given mRNA species and RNA-seq analyses of purified SGs revealed that signal sequence-encoding (i.e. endoplasmic reticulum (ER)-targeted) transcripts are significantly under-represented, consistent with prior reports that ER-localized mRNAs are excluded from SGs. Using translational profiling, cell fractionation, and single molecule mRNA imaging, we examined SG biogenesis during the unfolded protein response (UPR) and report that UPR-elicited SG formation is gene selective. Combined immunofluorescence-smFISH studies demonstrated that UPR-induced mRNA granules co-localized with SG protein markers and were in close physical proximity to or directly associated with the ER membrane. mRNA recruitment into ER-associated SGs required stress-induced translational inhibition, though translational inhibition was not solely predictive of mRNA accumulation in SGs. SG formation in response to UPR activation or arsenite addition was blocked by the transcriptional inhibitors actinomycin D or triptolide, suggesting a functional link between gene transcriptional state and SG biogenesis. These data demonstrate that ER-targeted mRNAs can be recruited into SGs and identify the ER as a subcellular site of SG assembly. On the basis of the transcriptional inhibitor studies, we propose that newly transcribed mRNAs undergoing nuclear export during conditions of suppressed translation initiation are key substrates for SG biogenesis.

## Introduction

Environmental, pathogen, and nutrient stressors can profoundly disrupt proteostasis, leading to toxic protein aggregation and in scenarios of unresolved stress, cell death (Harding et al. 2003; Sakaki et al. 2012; Wang and Kaufman 2012; Costa-Mattioli and Walter 2020). Reflecting the pathological consequences of dysregulated proteostasis, eukaryotic cells express a family of stress response eukaryotic initiation factor 2α (eIF2α) kinases whose activation results in the rapid inhibition of protein synthesis (Pakos-Zebrucka et al. 2016; Wek 2018). eIF2α kinase activity is central to the integrated stress response, which comprises a translational regulatory arm, mediated by the eIF2α kinases PERK, GCN2, PKR and HRI, and a stress response transcriptional program, whose activation supports the restoration of cellular proteostasis and promotes stress tolerance (Wek et al. 2006; Taniuchi et al. 2016).

In response to stress-induced activation of eIF2α kinase activity, translationally suppressed mRNAs can undergo recruitment into stress granules (SGs), membraneless organelles comprised of mRNAs and RNA binding proteins (RBPs) (Kedersha et al. 1999; Decker and Parker 2012; Wolozin and Ivanov 2019; Mateju et al. 2020). SG biogenesis is driven by the biochemical and biophysical properties of SG-resident RBPs, in particular multivalent RNA binding and intrinsically disordered domains, that in the presence of RNA support granule self-assembly (Kato et al. 2012; Guillen-Boixet et al. 2020; Matheny et al. 2020). mRNA recruitment into SGs is not, however, a simple process; the biochemical criteria for mRNA recruitment are complex and include intrinsic and stress-regulated translation efficiencies, transcript length, AU-rich element abundance, as well as other undefined *cis*-encoded elements (Khong et al. 2017; Namkoong et al. 2018; Matheny et al. 2020). In addition to these global criteria, deep sequencing analyses of SG RNAs revealed that endoplasmic reticulum (ER)-targeted transcripts are substantially under-represented (Khong et al. 2017). This finding is consistent with earlier reports that ER-targeted mRNAs are excluded from SGs (Unsworth et al. 2010). Intriguingly, the ER engages in dynamic interactions with SGs and processing bodies (PBs) which serve to regulate granule biogenesis and fission (Lee et al. 2020). How the ER contributes to SG and PB biology while its associated mRNAs are largely sequestered from these regulatory organelles is unknown.

As a physiological stress response pathway that regulates global translation initiation through eIF2α phosphorylation (Sidrauski et al. 2015), the unfolded protein response (UPR) provides a useful biological model for examining the functional interface between the ER and SG biology. In addition to its translation and gene regulatory arms described above, activation of the UPR elicits substantial subcellular mRNA localization dynamics, where ER-targeted mRNAs are released from the ER membrane and subsequently re-localized as the UPR progresses (Reid et al. 2014). Mechanistic links between RNA dynamics on the ER and SG biogenesis remain, however, largely unexplored.

Here we report that ER-targeted mRNAs can be recruited into SGs in response to DTT-elicited UPR activation or arsenite-induced oxidative stress in a gene-selective manner. Of the four ER-targeted mRNAs examined, SG accumulation was observed for HSP90B1/GRP94 and CTGF/CCN2, but not HSPA5/GRP78 or B2M. Selective SG recruitment was also observed for cytoplasmic mRNAs, where NCL was recruited into SG, but GAPDH was not. Combined cell fractionation and single molecule fluorescence *in situ* hybridization (smFISH) studies revealed that SGs formed at or on the ER membrane, identifying the ER as a subcellular site of SG biogenesis. Intriguingly, mRNAs experiencing similar translational suppression in response to UPR activation showed differential recruitment into SGs, indicating that translational suppression alone is insufficient for SG recruitment. SG formation was, however, highly sensitive to the transcriptional inhibitors ActD and triptolide, which prevented SG formation for all mRNAs examined. We summarize these data in a working model of SG assembly on the ER, emphasizing a role for newly exported mRNAs as preferred substrates for SG recruitment.

## Results

### Recruitment of ER-targeted mRNAs into stress granules

To examine the intersection between the ER, UPR, and ribonucleoprotein (RNP) granule dynamics, we first performed polyribosome profiling and single molecule RNA imaging of three ER-targeted mRNAs at steady state and following UPR induction. UPR activation was initiated by treatment of HeLa cells with the reducing agent 1,4-dithiothreitol (DTT) and monitored in assays of [^35^S]Met/Cys incorporation (**Fig. 1A**), eIF2α phosphorylation (**Fig. 1B**), and UPR-target gene transcriptional upregulation (**Fig. 1C**). DTT addition yielded a rapid reduction in cellular translation to ca. 20% of control at 30 min, followed by recovery to a reduced steady state (**Fig. 1A**). The biphasic translation inhibitory response was mirrored in the kinetics of eIF2α phosphorylation (**Fig. 1B**), where phospho-eIF2α levels peaked at 30-60 min of DTT treatment and gradually resolved after 120 min. Activation of the UPR transcriptional arm was assayed by RT-qPCR analysis of ER chaperone GRP78 and GRP94 mRNAs and revealed significant induction of both transcripts at 60 min, with near-maximal induction at 120 min (**Fig. 1C**). Levels of the ER-targeted B2M mRNA were not altered in response to DTT addition, consistent with prior studies demonstrating that B2M is not a UPR response gene (**Fig. 1C**). Sucrose density gradient polysome profiling studies confirmed global UPR-elicited translation inhibition as evidenced by the pronounced collapse of heavy polysomes (fractions 5-8, ribosome density > 4) to a predominately monosome profile (fractions 2-4, 80S monosome) following 60 min of DTT treatment (**Fig. 1D, E**, grey line). As with global polysome remodeling, UPR activation had a substantial impact on GRP94 and B2M translation profiles, with parallel shifts of their mRNAs from heavy to light polysome fractions indicating reduced translation initiation frequencies for these mRNAs (**Fig. 1D, E**). Notably, UPR activation had comparatively modest effects on theGRP78 translation profile, with a less pronounced redistribution of GRP78 mRNAs to light polysomes (**Fig. 1D, E**). The blunted translational response of GRP78 mRNAs to UPR activation is consistent with previous reports that its 5’ UTR encodes an internal ribosome entry site, enabling translation initiation under conditions of elevated eIF2α phosphorylation (Starck et al. 2016).

**Figure 1.**
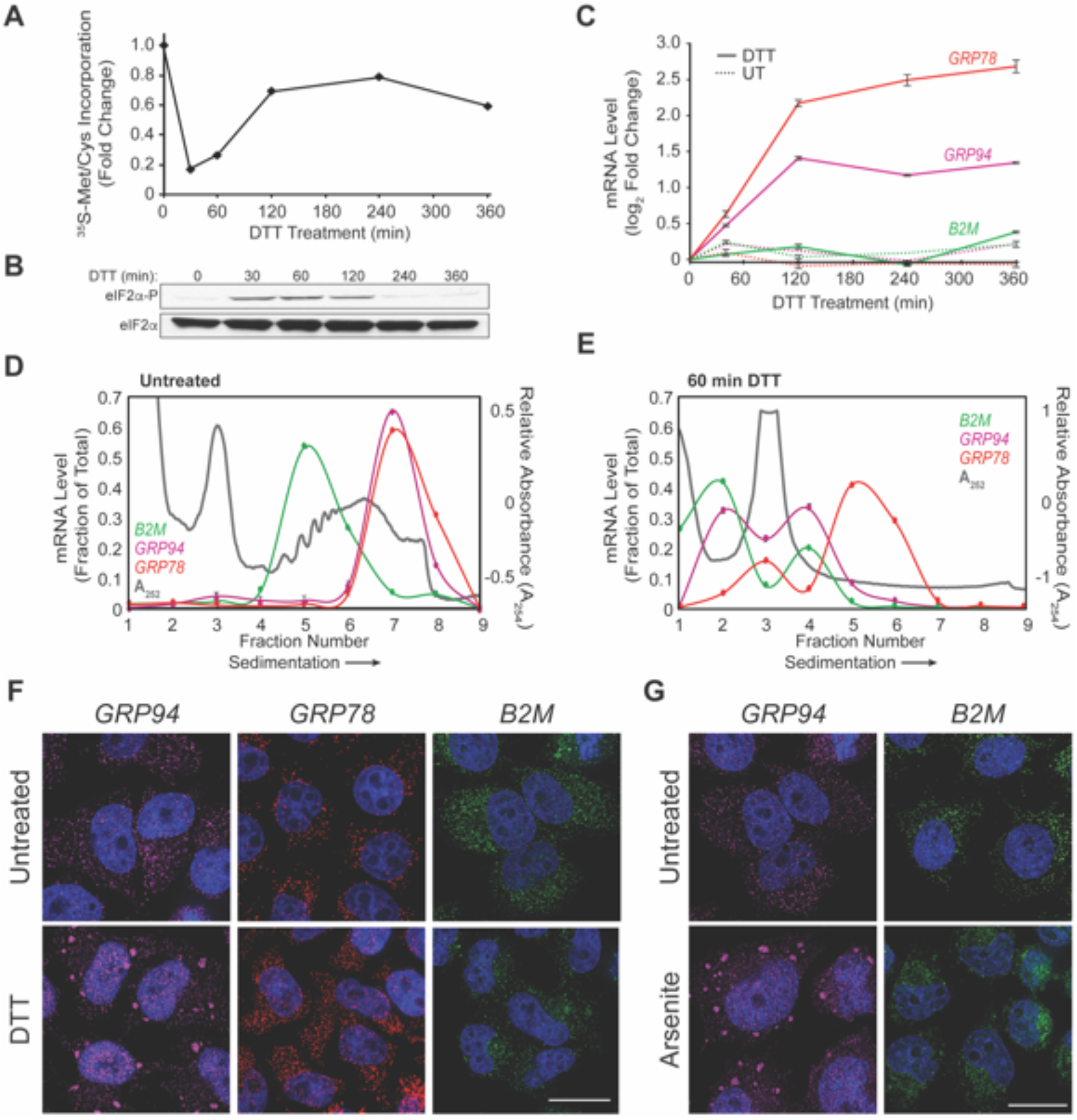
Selective recruitment of ER-targeted mRNAs into UPR-induced stress granules. (**A**) Time course of UPR-induced inhibition of protein synthesis assayed by [^35^S]Met/Cys incorporation. HeLa cell cultures were supplemented with DTT and protein synthesis rates assayed at the the indicated time points. (**B**) Immunoblot analysis of eIF2α phosphorylation in response to DTT addition. As in (**A**), cell cultures were supplemented with DTT and sampled for immunoblot analysis of eIF2α and phospho-eIF2α at the indicated time points. (**C**) Representative time course of UPR-elicited transcriptional activation of the UPR response genes GRP94 and GRP78 and the ER-targeted gene B2M. As in (**A**), cultures were supplemented with DTT and at the indicated time points, total RNA was extracted for RT-qPCR analysis of transcript levels. Data points are mean log_2_ fold-change ± SD, normalized to GAPDH levels. (**D**,**E**) Polyribosome profiling of GRP94, GRP78, and B2M translational status by sucrose density gradient velocity sedimentation. Hela cell cultures at time zero (**D**) or following DTT treatment (**E**) were detergent extracted and total polyribosome profiles and mRNA distributions were determined. Polyribosome patterns (grey) are represented by the A_254_ nm absorbance traces. mRNA distributions were determined by RT-qPCR analysis of GRP78 (red), GRP94 (magenta), and B2M (green) mRNAs. Data are representative of three biological replicates; RT-qPCR data are mean fraction of total mRNA for the given gene throughout all gradient fractions ± SD. (**F**) Representative images of smFISH visualization of GRP94 (magenta), GRP78 (red), and B2M (green) mRNAs in untreated and DTT-treated (60 min) HeLa cells. (**G**) As in (**F**) but treatment with sodium arsenite (60 min). DAPI nuclear stain (blue) is indicated for all images. Scale bar = 20 μm.

With translation initiation inhibition being a primary trigger for SG formation (Kedersha et al. 1999), we examined the subcellular distributions of GRP94, GRP78, and B2M mRNAs by single molecule fluorescence *in situ* hybridization (smFISH) before and after UPR activation (**Fig. 1F**). For the two mRNAs whose translation was strongly repressed in response to UPR activation, i.e. GRP94 and B2M, divergent smFISH patterns were observed. GRP94 mRNAs transitioned from diffuse, diffraction-limited single foci at steady state to accumulated prominent perinuclear granules following UPR activation, whereas B2M mRNAs were maintained as single foci in both untreated and UPR-activated conditions. As expected for actively translating mRNAs, GRP78 smFISH patterns were largely unaltered by the UPR (**Fig. 1F**). These data indicate that mRNA recruitment into UPR-elicited granules is not solely driven by mRNA translational status, and suggest that gene-specific phenomenon may contribute to granule formation. Importantly, these findings were not unique to UPR stress as treatment of HeLa cell cultures with the oxidative stressor sodium arsenite also elicited GRP94, but not B2M, mRNA granule formation (**Fig. 1G**).

### Stress granules containing ER-targeted mRNAs can be ER-associated

With prior studies reporting that ER-targeted mRNAs were under-represented and/or excluded from SGs (Unsworth et al. 2010; Khong et al. 2017), we considered that the UPR-elicited GRP94 mRNA granules could comprise a novel RNP granule, distinct from canonical SGs. To examine this hypothesis, immunofluorescence co-localization studies were performed for the SG marker proteins HuR, G3BP1, and PABP in parallel with GRP94 smFISH (Stoecklin and Kedersha 2013; Protter and Parker 2016; Youn et al. 2018). GRP94 mRNA granules co-stained with the three SG components following but not prior to UPR activation, consistent with recruitment of GRP94 mRNAs into canonical, rather than unique, SGs (**Fig. 2A**).

**Figure 2.**
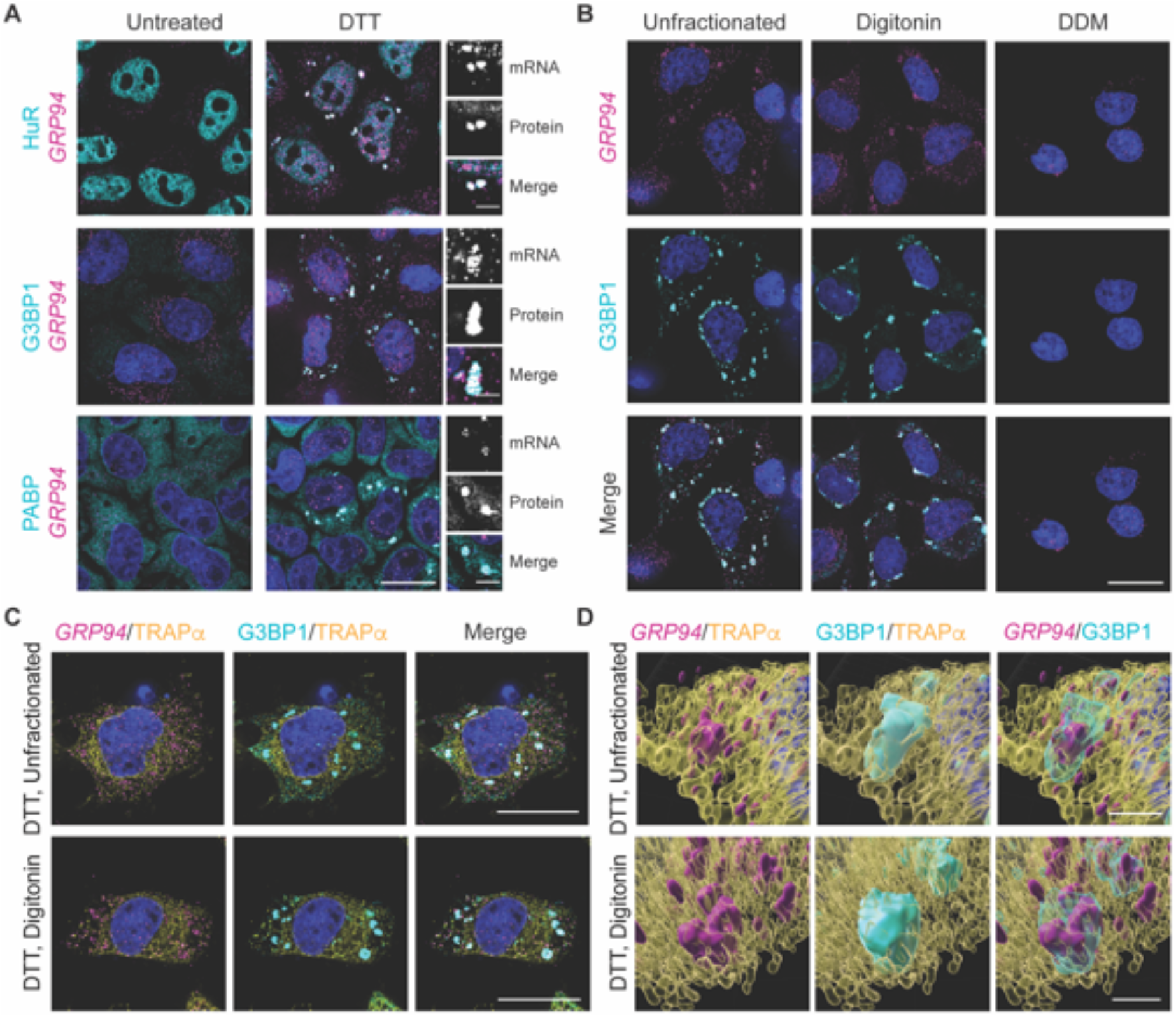
UPR activation elicits stress granule formation on the ER membrane. (**A**) Representative GRP94 smFISH (magenta) with immunofluorescence co-staining for the stress granule protein markers HuR, G3BP1, and PABP (cyan) in untreated and DTT-treated HeLa cell cultures. Grayscale insets for mRNA and protein channels, as well as color merge, for DTT-treated cells (right). (**B**) Representative GRP94 smFISH (magenta) and G3BP1 immunofluorescence (cyan) co-staining in DTT-treated cells. Following DTT treatment, cells were permeabilized with digitonin-supplemented buffers to release cytosolic contents (digitonin) or sequentially treated with digitonin and n-dodecyl-β-D-maltoside (DDM) buffers to solubilize organelle membranes. (**C**) Representative micrographs of ER membrane protein TRAPα immunofluorescence (yellow) with GRP94 smFISH (magenta) and/or G3BP1immunofluorescence (cyan) co-staining in unfractionated and cytosol-depleted (digitonin-permeabilized) cells following DTT treatment. (**D**) 3D renderings of representative granules from (**C**) in unfractionated (**Movie 1**) and digitonin permeabilized (**Movie 2**) cells. DAPI staining (blue) is indicated for all images. Full cell scale bars = 20 μm, inset and 3D scale bars = 4 μm.

With GRP94 mRNAs displaying high enrichment on the ER (Chen et al. 2011; Reid and Nicchitta 2012; Reid et al. 2014), we asked if GRP94-containing SGs were also ER-associated. To examine the subcellular distribution of GRP94 SGs, HeLa cell cultures were treated with DTT and GRP94 smFISH patterns were compared between intact, cytosol-depleted, and cytosol/ER-depleted cells using a sequential detergent fractionation protocol that efficiently separates the cytosol and membrane compartments of tissue culture cells (**Fig. 2B, Fig. S1**) (Lerner et al. 2003; Jagannathan et al. 2011). Here, the subcellular distribution patterns of the GRP94 SGs and associated G3BP1 protein were retained following the digitonin-dependent release of cytosolic cellular contents. Subsequent treatment with the ER membrane-solubilizing detergent dodecylmaltoside (DDM) resulted in the complete loss of extranuclear GRP94 smFISH signal and G3BP1 immunostaining, indicating that GRP94 SGs were membrane-associated (**Fig. 2B**). The minor intranuclear GRP94 smFISH signal retained following DDM treatment demonstrates a selective loss of GRP94 SGs coincident with ER membrane solubilization while the nucleus remains largely intact.

To further explore the ER-association of SGs, GRP94 smFISH was performed with co-immunostaining for the ER-resident membrane protein TRAPα and the SG marker G3BP1 in unfractionated and cytosol-depleted cells (**Fig. 2C**). The retention of coincident GRP94 mRNA and G3BP1 staining in cytosol-depleted cells supports the conclusion that SG formation occurred in close physical proximity or in direct association with the ER membrane. 3D-reconstruction of these images (**Movie 1, Movie 2**) were consistent with this conclusion and revealed SG complexes engaged in apparent contact sites with the ER (**Fig. 2D**). By the criteria of detergent sensitivity and fluorescence co-localization, these data demonstrate that ER-targeted mRNAs can be recruited to ER-associated SGs and identify the ER as a subcellular site of SG biogenesis (Khong et al. 2017; Lee et al. 2020).

### Transcriptional inhibitors block GRP94 stress granule formation

The data in **Fig. 1F** indicate that mRNA recruitment into ER-associated SGs was selective for one of the three mRNAs examined, GRP94. mRNAs can differ in their translational sensitivity to eIF2α phosphorylation state and thus a time course study was performed to examine SG formation over a time period that captures both the upregulation and resolution phase of eIF2α phosphorylation during DTT treatment (**Fig. 1A, B**) (Andreev et al. 2015; Sidrauski et al. 2015; Young and Wek 2016). In these experiments, GRP94 granules were observed as early as 30 min post-DTT addition whereas B2M mRNAs remained as diffuse single foci throughout the two hour time course (**Fig. 3A**) despite the similar and sustained inhibition of translation for the two mRNAs (**Fig. 3B**). The smFISH data depicted in **Fig. 3A** further highlight the divergent transcriptional responses of the two genes to UPR activation. For GRP94, the UPR-elicited transcriptional induction was detected by smFISH as intranuclear transcriptional foci at the 30 min time point, with prominent intranuclear mRNA staining at the 120 min time point (**Fig. 3A**). In contrast, UPR activation did not alter B2M intranuclear smFISH patterns, consistent with the data presented in **Fig.1B2**. These smFISH data provide orthogonal validation that GRP94, but not B2M, is transcriptionally upregulated in response to UPR activation and indicate that gene transcriptional state may be a criterion for mRNA recruitment into SGs.

**Figure 3.**
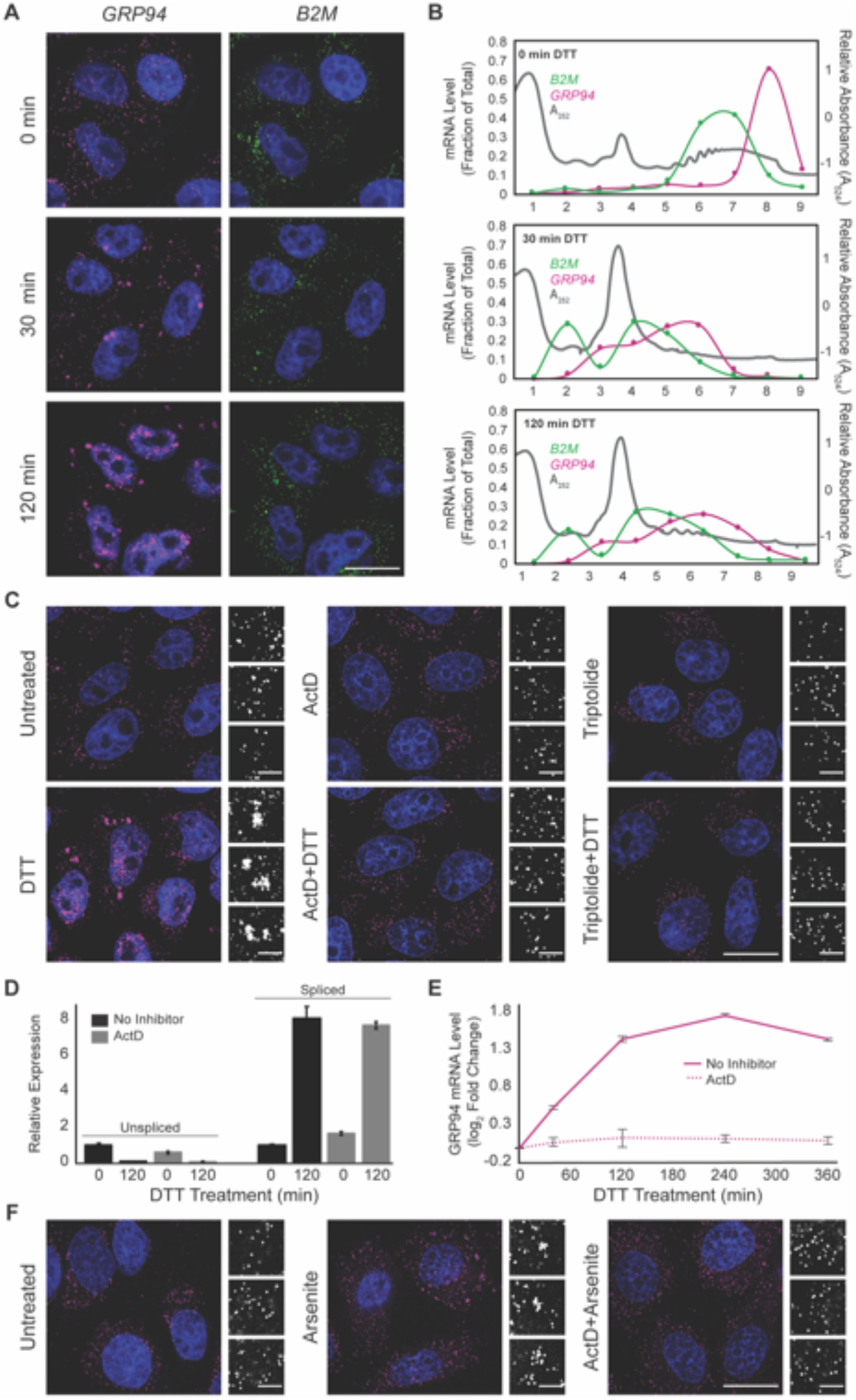
Transcription inhibitors ActD and triptolide prevent stress granule formation. (**A**) Representative time course analysis of GRP94 (magenta) and B2M (green) smFISH staining patterns over the time course of maximal inhibition of eIF2α activity (see Fig.1A). Note appearance of intranuclear transcriptional hotspots and mRNA accumulation in the GRP94 smFISH analyses, absent in the B2M smFISH staining. (**B**) Sucrose density velocity sedimentation gradients and RT-qPCR analysis as in **Fig. 1D** at the indicated time points following DTT addition. Data is representative of three biological replicates. (**C**) Representative GRP94 smFISH in control (untreated, ActD, triptolide) and stressed (treatment with DTT, with and without treatment with indicated transcriptional inhibitor, ActD or triptolide) conditions. Grayscale insets of mRNA distributions are provided for each condition (right). (**D**) RT-qPCR analysis of spliced and unspliced XBP1 mRNA in control (untreated or ActD) and stressed (treatment with DTT, with and without ActD) conditions. Expression levels at 0 min without ActD were set as 1 after normalization to GAPDH. *n* = 2 biological replicates. Data are means ± SD. (**E**) RT-qPCR analysis of GRP94 mRNA levels over a time course of DTT treatment with or without ActD addition. *n* = 2 biological replicates. Data are mean log_2_ fold change of expression relative to GAPDH ± SD. (**F**) Representative images GRP94 smFISH in untreated and sodium arsenite-treated cells with or without ActD addition. Grayscale insets of mRNA distribution are provided for each condition (right). DAPI staining (blue) included in all images. Full cell scale bar = 20 μm, inset scale bar = 4 μm.

To explore potential links between gene transcription and mRNA recruitment into SGs, we performed GRP94 smFISH studies in cells treated with the transcriptional inhibitors actinomycin D (ActD), a DNA intercalating agent, or triptolide, an RNA polymerase II inhibitor (Titov et al. 2011). As shown in **Fig. 3C**, DTT-elicited GRP94 SG formation was efficiently inhibited in the presence of either ActD or triptolide. To exclude possible off-target effects on UPR signaling, we assayed for IRE1-mediated splicing of transcription factor XBP1 mRNA, a signature event in UPR activation, in the absence or presence of ActD. As expected, ActD addition resulted in decreased levels of unspliced XBP1 mRNA at the two hour time point, a consequence of mRNA decay in the absence of ongoing transcription. ActD did not, however, alter IRE1 activation, demonstrated by similar levels of spliced XBP1 mRNA in both control and ActD-treated cells following DTT addition (**Fig. 3D**). These data demonstrate that transcriptional inhibition did not disrupt UPR signaling, however ActD treatment did prevent transcriptional upregulation of GRP94 (**Fig. 3E**). To determine if the connection between active transcription and SG biogenesis was unique to UPR-elicited SGs, we assayed arsenite-stimulated SG assembly under identical conditions and observed that ActD treatment prevented GRP94 SG formation in response to arsenite addition (**Fig. 3F**).

As with arsenite-induced SGs, GRP94 SGs contain G3BP1, PABP, and HuR (**Fig. 2A**), suggesting they may share overlapping mechanisms of formation. To explore this further, we investigated the effects of ActD treatment on the recruitment of HuR and PABP into SGs (**Fig. 4**). Notably, ActD treatment blocked formation of HuR- and PABP-positive granules following DTT addition (**Fig. 4A, B**). As reported previously, ActD treatment evoked the redistribution of HuR from the nucleoplasm to the cytoplasm, without altering the subcellular distribution of PABP, and this phenomenon was not affected by DTT addition (Peng et al. 1998; Bounedjah et al. 2014). Consistent with the view that UPR-stimulated SG formation occurs through processes similar to arsenite-stimulated SG formation, treatment with the protein synthesis inhibitor cycloheximide blocked PABP and GRP94 SG formation elicited by either DTT (**Fig. 4C**) or arsenite (**Fig. 4D**). Combined, these data reveal that transcriptional inhibition prevents GRP94 SG formation and suggest a role for *de novo* RNA transcription in SG biogenesis.

**Figure 4.**
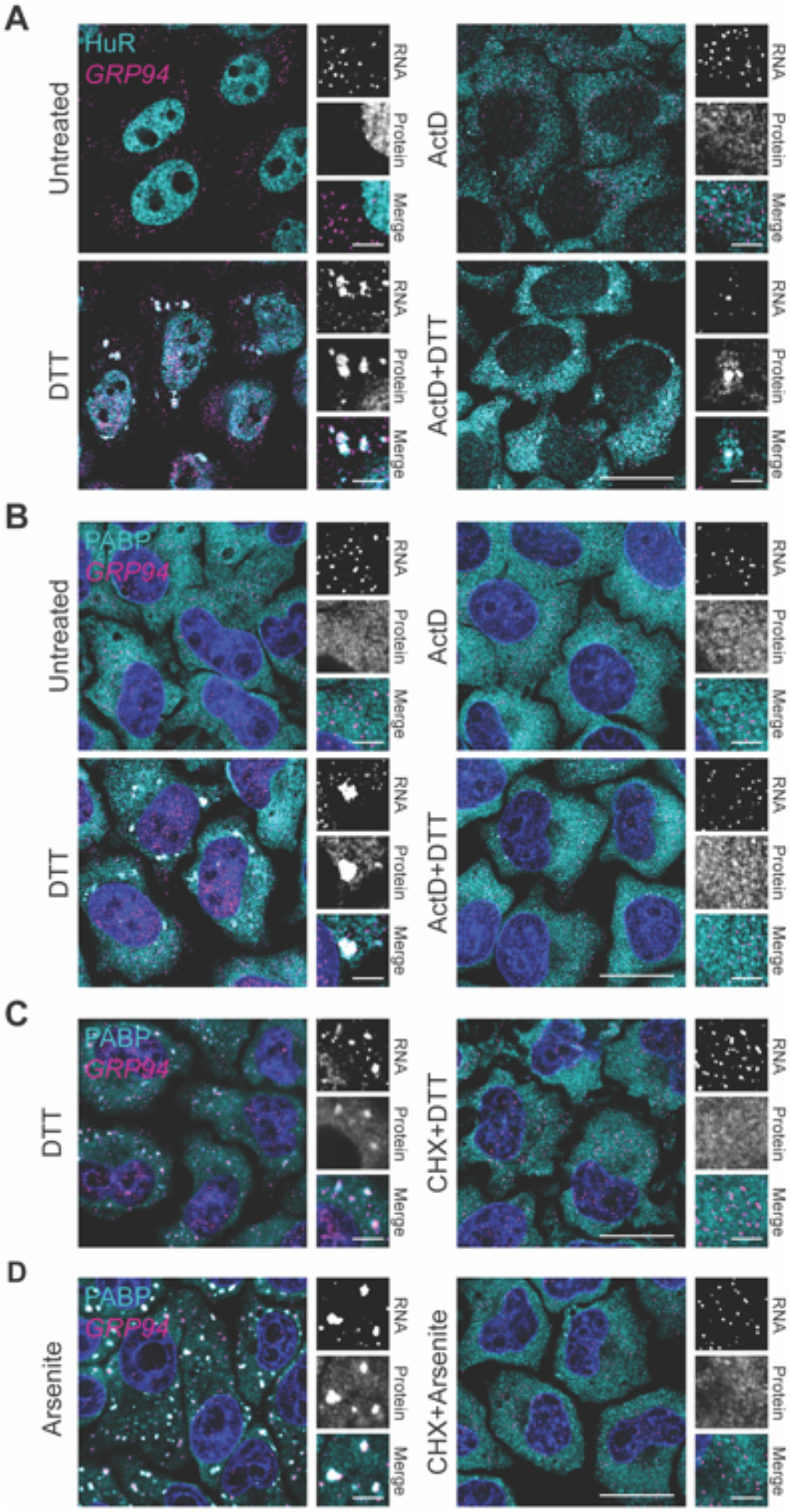
Actinomycin D prevents RNA binding protein recruitment into UPR-elicited stress granules. (**A**) Representative GRP94 smFISH (magenta) with immunofluorescence co-staining for HuR (cyan) in control (untreated or ActD) and stressed (treatment with DTT, with and without ActD) conditions in HeLa cell cultures. Note redistribution of HuR from the nucleoplasm to the cytoplasm in response to ActD treatment. Grayscale insets for mRNA and protein channels, as well as color merge images, for each condition are provided (right). (**B**) As in (**A**) but immunofluorescence staining for PABP (cyan). (**C**) Representative GRP94 smFISH (magenta) with immunofluorescence co-staining for PABP (cyan) in cells treated with DTT treatment with and without cycloheximide (CHX) addition. (**D**) As in (**C**) but with sodium arsenite stress. DAPI nuclear stain (blue) is included in all images. Full cell scale bar = 20 μm, inset scale bar = 4 μm.

### Transcriptional inhibition suppresses stress granule recruitment of cytoplasmic and ER-targeted mRNAs

The data presented in **Fig. 3** and **4** indicate that pharmacological inhibition of transcription disrupts ER-associated SG formation. We also considered that this phenomenon may be restricted to ER-targeted mRNAs and/or ER-associated SG biogenesis. To further explore the association between ER-targeting and SG recruitment sensitivity to transcriptional inhibitors, we selected additional genes for smFISH studies of SG biogenesis. With SG recruitment efficiencies positively correlating with CDS length (Khong et al. 2017), transcript length was also considered during gene selection. Using these criteria, three genes were chosen: nucleolin, connective tissue growth factor (CTGF/CCN2), and glyceraldehyde 3-phosphate dehydrogenase (GAPDH). NCL (CDS = 2133 nt) encodes a transcript of similar length to GRP94 (CDS = 2412 nt) and lacks an encoded signal sequence, therefore identifying it as a cytoplasmic mRNA, allowing for assessment of the requirement for ER-targeting in recruitment into ER-associated SG. CCN2 (CDS = 1050 nt) was previously identified as an ER-targeted UPR-responsive gene (Rendleman et al. 2018), but encodes a substantially shorter transcript than GRP94, providing a useful test of the contribution of gene transcriptional status vs. transcript length to SG recruitment among ER-targeted genes (Khong et al. 2017). GAPDH (CDS = 1008 nt) was selected as a length-matched comparator for CCN2 that, like NCL, lacks an encoded signal sequence and whose transcription is not UPR-responsive (Rendleman et al. 2018).

Paired smFISH of GRP94, NCL, CCN2, and GAPDH mRNAs were performed at steady state and following UPR activation with and without transcriptional inhibition. As depicted in **Fig. 5**, GRP94, NCL, and CCN2 mRNAs were recruited to perinuclear SGs following DTT addition, whereas GAPDH mRNAs displayed a diffuse subcellular distribution pattern very similar to the untreated control. Consistent with data reported in **Fig. 3**, treatment of cell cultures with either ActD or triptolide blocked UPR-induced SG formation for GRP94, NCL, and CCN2 alike, further suggesting a functional link between newly transcribed/exported mRNAs and SG biogenesis. To determine if UPR-induced NCL and CCN2 SGs were ER-associated, as was observed for GRP94 SGs (**Fig. 2**), cells were treated with DTT and mRNA distributions examined following digitonin extraction of the cytosol (DTT + digitonin) (**Fig. 5**). These data demonstrate that NCL and CCN2 granules were also retained in perinuclear locales in digitonin-permeabilized cells, consistent with an ER association. Combined, these data support a model where SG biogenesis can occur on or in immediate physical proximity to the ER regardless of an encoded signal sequence, and is disabled following transcriptional inhibition. These data are also consistent with the prior finding that transcript length is positively correlated with SG recruitment, where longer transcripts (e.g. GRP94 and NCL) were recruited to SGs and short transcripts (e.g., GAPDH, B2M) were refractory to SG recruitment (Khong et al. 2017). CCN2, as a short transcript that did assemble into SGs, provides an interesting case which suggests transcriptional activation may outweigh the transcript length predictions for mRNA recruitment into SGs. As discussed below, these data also confirm prior studies reporting a broad representation of the mRNA transcriptome on the ER (Diehn et al. 2000; Lerner et al. 2003; Lerner and Nicchitta 2006; Chen et al. 2011; Reid and Nicchitta 2012; Jan et al. 2014; Reid and Nicchitta 2015a; Chartron et al. 2016).

**Figure 5.**
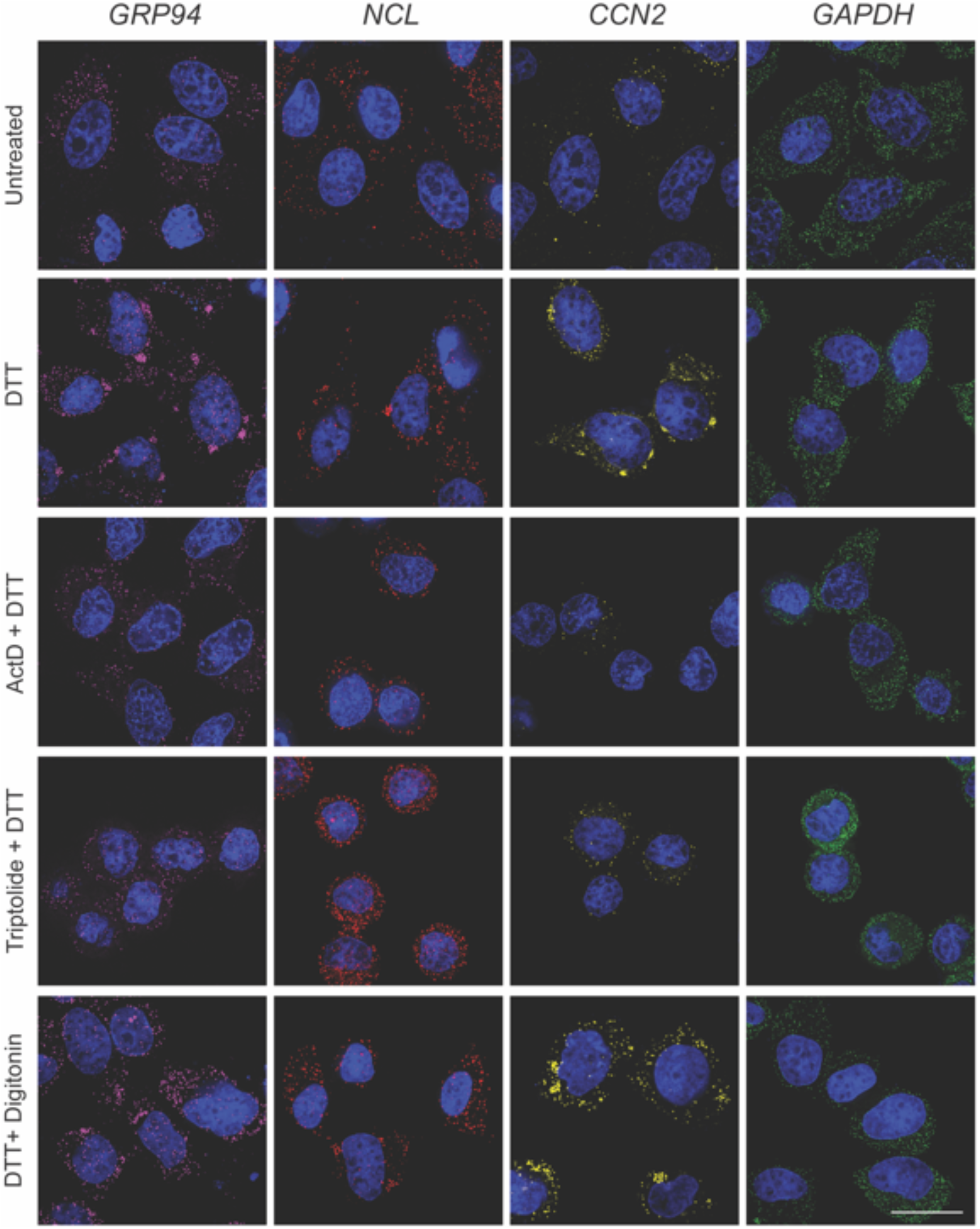
Transcript length and transcriptional activation state influence mRNA recruitment into ER-associated stress granules. (**A**) Representative micrographs of long transcript size-paired genes GRP94 (magenta, 2412 nt) and NCL (red, 2133 nt), and short transcript size-paired genes CCN2 (yellow, 1050 nt) and GAPDH (green, 1008 ntbp) smFISH in untreated and DTT-treated HeLa cell cultures. Where indicated, cell cultures were treated with ActD or triptolide in addition to DTT. To distinguish between cytosolic and ER-associated SGs, cells were permeabilized in digitonin-supplemented buffers to extract cytosol contents, while leaving the ER membrane intact. DAPI nuclear stain (blue) included in all images. Scale bar = 20 μm.

## Discussion

Activation of the UPR, an ER-centric stress response signaling pathway, causes dramatic remodeling of translation on the ER (Stephens and Nicchitta 2008; Reid et al. 2014; Reid and Nicchitta 2015a). Via the UPR sensor/eIF2α kinase, PERK, translation initiation is reduced following UPR activation, and stress granule formation is enabled (Namkoong et al. 2018; Reich et al. 2020). Notably, however, ER-targeted mRNAs are underrepresented in SG transcriptome analyses and can be excluded from SGs (Unsworth et al. 2010; Khong et al. 2017). Here we report examples of ER-targeted mRNAs that are efficiently recruited into SGs in response to both UPR activation and arsenite stress. With the constraint of a limited number of mRNAs examined, this phenomenon was found to be gene-specific, where the signal sequence-encoding mRNAs GRP94/HSP90B1 and CCN2/CTGF were recruited into SGs whereas GRP78/BiP/HSPA5 and B2M were refractory to SG recruitment. Unexpectedly, GRP94 and CCN2 SGs were found to form in close apposition to, or in direct association with, the ER membrane, suggesting the ER may serve as an important subcellular locale for SG biogenesis. Although little is currently known regarding the subcellular organization of SG dynamics, the recent finding that ER contact sites regulate granule formation and fission support a role for the ER in SG biology. We also report that inhibition of transcription with either the DNA intercalating agent, ActD, or the RNA polymerase II inhibitor, triptolide, blocked UPR- and arsenite-elicited SG formation for both ER-targeted and cytoplasmic mRNAs. The sensitivity of SG biogenesis to the presence of transcriptional inhibitors suggests that gene transcriptional state may be a relevant criterion in the complex process of SG assembly.

The finding that ER-targeted mRNAs vary in their SG recruitment patterns indicates that, like translational status, an encoded signal sequence is not itself prognostic of mRNA recruitment into SGs. The conclusion that select ER-targeted mRNAs can assemble into SGs was somewhat unexpected, given that ER-targeted mRNAs are under-represented in SG transcriptomes and that ER markers are nearly absent in purified SGs (Khong et al. 2017; Namkoong et al. 2018). Additionally, a prior analysis of the translational and SG recruitment response of the ER-localized mRNA MDR1 to arsenite stress demonstrated that MDR1 transcripts are entirely refractory to SG recruitment (Unsworth et al. 2010). In the absence of a detailed understanding of the criteria for mRNA recruitment into SGs, it’s difficult to conclude that these findings are necessarily incongruent. For example, in the case of SG transcriptome analyses, the biochemical purification methods used for SG isolation may not yield a fully representative SG population, as noted by the authors (Khong et al. 2017). Another potentially important consideration regarding the purified SG transcriptome is that, at the mRNA level, ER-targeted genes, which largely encode secreted or integral membrane proteins, are expressed at relatively low levels and are more cell-type variable compared to housekeeping genes (Reid and Nicchitta 2012; Reid and Nicchitta 2015a; Reid and Nicchitta 2015b). These factors may preclude detection of ER-targeted genes from purified subsets of RNA populations in a given cell type. Transcript abundance itself, however, is unlikely to be a primary driver for SG recruitment as is evident by experiments with GAPDH mRNAs (**Fig. 5**), which, although highly abundant, did not undergo SG assembly in the stress conditions examined here. Lastly, bioinformatic analyses of the SG transcriptome have identified a positive correlation between SG recruitment efficiency and transcript length. This positive correlation was supported by the mRNAs examined in the current study; longer transcripts such as GRP94 and NCL mRNAs were recruited into SGs whereas shorter transcripts such as B2M and GAPDH were refractory to SG recruitment (Khong et al. 2017; Namkoong et al. 2018). Exceptions to this length-based correlation are also apparent in that MDR1 encodes long transcripts (CDS: 3843 nt), are ER-associated, and yet escape SG recruitment (Unsworth et al. 2010), and CCN2 transcripts reported within are relatively short, ER-associated, and are efficiently recruited into SGs. One element that could explain these exceptions is UPR-induced transcriptional upregulation; GRP94 and CCN2 mRNAs are UPR responsive, whereas MDR1 is not (Rendleman et al. 2018). This hypothesis is supported by the current finding that transcriptional inhibition prevents SG formation.

Although substantial progress has been made in defining the protein and RNA compositions of SGs, as well as their exchange dynamics, little is known regarding their subcellular localization. The discovery of ER contact sites that regulate RNP granule formation and fission is consistent with the view that the ER may contribute to SG biology (Lee et al. 2020). To this point, the mRNA transcriptome and translatome is broadly represented on the ER membrane, supporting the observation of both ER-targeted and cytoplasmic mRNAs being recruited into ER-associated SGs (Diehn et al. 2000; Chen et al. 2011; Reid and Nicchitta 2012; Jan et al. 2014; Reid and Nicchitta 2015a; Chartron et al. 2016; Voigt et al. 2017). Furthermore, mRNAs can undergo direct (i.e., ribosome-independent) anchoring to the ER and comprehensive proteomic screens for RNA-interacting proteins have identified a number of candidate ER integral membrane RBPs that, by virtue of their ability to localize mRNAs to the cytoplasmic surface of the ER, may assist in ER-associated SG biogenesis (Cui et al. 2012; Jagannathan et al. 2014; Castello et al. 2016; Hsu et al. 2018; Queiroz et al. 2019; Trendel et al. 2019). Notably, one of these proteins (AEG1/MTDH) interacts with both ER-targeted and cytoplasmic-encoding mRNAs and, although lacking known RNA binding motifs, contains a large intrinsically disordered domain, a frequently identified motif thought to function in SG nucleation (Hsu et al. 2018). These observations suggest that insights into the subcellular locale(s) of SGs, their dynamic interactions with intracellular membranes, and the biochemical mechanism of membrane tethering may represent important avenues of future study.

In consideration of the criteria which may predict mRNA recruitment into SGs, the identification of two ER-targeted mRNAs whose transcription is upregulated in response to UPR activation and which undergo recruitment into SGs prompted us to ask if *de novo* transcription may contribute to SG biogenesis. Using two inhibitors of transcription with different modes of action, as well as two separate stress conditions, we observed transcriptional inhibition efficiently blocked SG formation. In the scenario of the UPR, transcriptional inhibition did not disrupt activation of the stress sensor/endonuclease IRE1, assayed as production of the UPR transcription factor XBP1s, indicating that the lack of SG formation under these conditions was either a direct or indirect consequence of the transcriptional block. Mechanistic links between gene transcriptional status and SG recruitment have not been widely investigated. However, a prior study examining SG formation in the context of poliovirus infection reported that ActD treatment blocked poliovirus infection-dependent SG formation (Piotrowska et al. 2010). These authors also observed that ActD treatment reduced SG formation, assayed via oligo-dT FISH, in response to arsenite and heat shock stress, and suggested that newly synthesized and preexisting mRNAs undergo separate and selective modes of recruitment into SGs. Bridging from this report, and noting that transcriptional upregulation is an integral element of the UPR, the finding that transcriptional inhibition blocks mRNA recruitment into ER-associated SGs led us to favor the view that newly exported mRNAs are a preferred substrate for SG assembly. The link to UPR activation also suggests that transcriptional status may be relevant to SG recruitment, where genes undergoing relatively high transcriptional rates, either via basal transcription or through UPR-activated transcription, represent high probability candidates for SG recruitment. Although the majority of our experiments involve UPR activation, which includes a transcriptional regulatory arm, it should also be noted that arsenite addition also evokes substantial changes in the cellular transcriptional program (Li et al. 2011; Jacobson et al. 2012; Srivastava et al. 2013; Delaney et al. 2020). Therefore, understanding how transcriptional upregulation impacts granule biogenesis may be of broad relevance to SG biology.

This perspective is summarized in a working model that considers existing links between transcription, nuclear export, translation-driven mRNP remodeling, and the subcellular trafficking fate of exported mRNAs in the context of SG biogenesis (**Fig. 6**) (Moore 2005; Palazzo et al. 2007; Martin and Ephrussi 2009; Maquat et al. 2010; Blower 2013). As illustrated, under steady-state conditions, newly exported mRNPs undergo efficient translation-coupled RBP remodeling coincident with export and are then recruited into the polyribosome-engaged mRNA pool (Lejeune et al. 2002; Maquat et al. 2010; Trcek et al. 2013; Halstead et al. 2015; He and Jacobson 2015). In contrast, under conditions of repressed translation initiation, newly exported mRNAs are inefficiently translated and thus retain a nuclear RBP signature for extended time periods post-export, potentially marking this population of mRNAs for SG recruitment (Dostie and Dreyfuss 2002; Maquat et al. 2010; Matheny et al. 2020). As previously reported, CDS and overall transcript length positively contribute to this process, perhaps through proportionately higher RBP binding densities per transcript and enhanced sterically-driven RNA-RNA interactions (Matherly et al. 1989; Namkoong et al. 2018; Hofmann et al. 2021). Following recovery from stress, translation-driven RBP remodeling is predicted to then confer resistance to subsequent SG recruitment (Piotrowska et al. 2010). By this mechanism, SGs may serve as storage depots for newly exported transcripts during stress, in particular for mRNAs comprising stress response gene programs, and subsequently as sites for translation-driven mobilization of these transcripts into the ribosome-engaged mRNA pool during recovery to cellular homeostasis. With the ER being physically continuous with the outer nuclear envelope, physical proximity may favor recruitment of newly exported mRNAs into ER-associated SGs, a view supported by recent work implicating the ER in SG and PB biogenesis (Lee et al. 2020).

**Figure 6.**
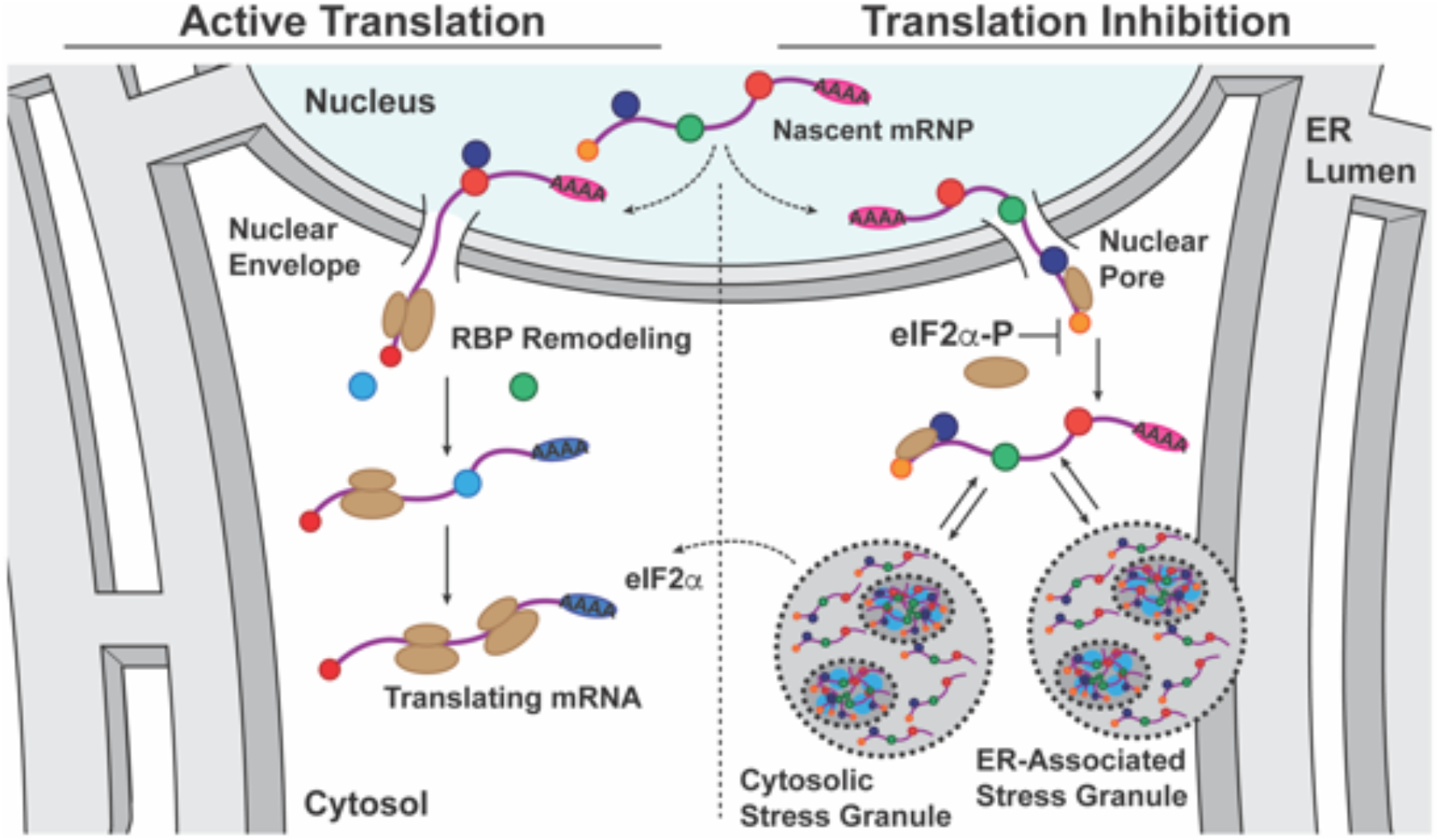
**Model depicting partitioning of newly exported mRNAs between translation-engaged and stress granule-associated states in response to stress pathway activation**. A working model depicting the functional states of newly exported mRNAs in homeostatic (active translation, low phospo-eIF2α) or stress-activated (reduced translation, elevated phospo-eIF2α) cellular states. This model highlights a role for translation-dependent RNA binding protein remodeling of newly exported mRNAs in determining mRNA recruitment into polyribosomes or SGs. As illustrated, under conditions of stress-induced translational inhibition, the nuclear RNA binding protein signature of newly exported mRNAs would be relatively long-lived and thus serve as a signal for ribonucleoprotein recruitment into stress granules, which could include both cytoplasmic and ER-associated forms. Once eIF2α phosphorylation is resolved, stress granule-associated mRNAs would resume translation, undergo RNA binding protein remodeling, and enter the polyribosome-associated pool. In this way, stress granules could serve as a triage station for newly exported mRNAs and undergo rapid mobilization following stress adaptation.

## Materials and Methods

### Cell culture and treatments

HeLa cells were cultured in DMEM plus 10% FBS at 37°C in 5% CO_2_ and experiments performed with cultures at 70-80% confluence. For experiments involving optical imaging, cells were seeded onto 12 mm round #1 glass coverslips. Polylysine (Sigma, Cat. No. P8920) coated glass coverslips were used for all optical imaging experiments where cells were subjected to sequential detergent fractionation prior to paraformaldehyde fixation. Where indicated, cell stress was elicited by treating cell cultures with 1-5 mM DTT or 1 mM arsenite for the noted time period (0-180 min). All DTT treatment times and concentrations were titrated for each batch of cells and DTT stock to ensure phenotypic consistency. For RNA transcription inhibitor experiments, cell cultures were treated with 10 μM actinomycin D or 30 nM triptolide for 30 min prior to stress induction and inhibitors were included in all media through the experimental time course. For translation inhibitor experiments, cell cultures were treated with 100 μg/mL cycloheximide for 10 min prior to stress induction and cycloheximide was included in all media through the experimental time course.

### [^35^S]-Met/Cys biosynthetic labeling

For determination of protein synthesis rates under conditions of UPR activation, cell cultures were treated with DTT for 0-6 hours and pulse-labeled for 10 min at the indicated time points with 75 μCi/mL [^35^S]-Met/Cys in methionine- and cysteine-free culture media. At the termination of the pulse labeling period, cell cultures were washed with ice-cold PBS containing 100 μg/mL cycloheximide to block further translation and subsequently lysed on ice for 5 min in 1% Triton X-100/PBS containing 100 μg/mL cycloheximide and protease inhibitor cocktail. Detergent lysates were adjusted to 10% (vol:vol) trichloroacetic acid, samples were incubated on ice, and precipitated proteins were recovered by centrifugation (15 min, 15,000 x *g*, 4°C). Protein pellets were washed with 100% acetone to remove residual TCA, allowed to air dry, and solubilized by addition of 0.5 M Tris, pH 11, 5% SDS. [^35^S]-Met/Cys levels were determined by liquid scintillation counting of the resuspended protein fraction and corresponding protein concentrations were determined by BCA assay (Pierce). Data are reported as scintillation counts per minute (CPM) per milligram (mg) of protein..

### Immunoblotting

For immunoblot analysis, HeLa cell cultures were placed on ice and lysed in ice-cold 1% CHAPSO, 150 mM NaCl, 20 mM K-HEPES, pH 7.5, 10% glycerol, 1 mM EDTA, 100 mM NaF, 17.5 mM B-glycerophosphate, and 1X protease inhibitor cocktail for 5 min. Lysates were TCA precipitated as noted above, washed with 100% acetone, resuspended in 0.5 M Tris, pH 11, 5% SDS, and protein concentrations determined via BCA assay (Pierce). Equivalent protein mass per sample was separated by SDS-PAGE, transferred to nitrocellulose membranes, blocked in 10% nonfat dry milk/Tris-buffered saline and processed per antibody supplier’s recommendations. Antibodies used include eIF2α (Santa Cruz, Cat. No. sc11386; 1:500 dilution) and phospho-eIF2α (Cell Signaling Technologies, Cat. No. 3398; 1:1500 dilution).

### Polysome gradients, RNA extraction, and RT-qPCR analyses

Polyribosomes from cells lysed in dodecylmaltoside (DDM) lysis buffer (200 mM KCl, 25 mM K-HEPES, pH 7.2, 10 mM MgOAc_2_, 2 mM DTT, 50 μg/mL cycloheximide, 1X protease inhibitor cocktail, 40U/mL RNaseOUT, and 2% DDM) were resolved on 15-50% sucrose gradients and fractionated using a Teledyne/Isco gradient fractionator as described in Stephens and Nicchitta (Stephens and Nicchitta 2007; Stephens and Nicchitta 2008). Total RNA was extracted from collected gradient fractions or total cell lysates by guanidinium thiocyanate-phenol-chloroform extraction and RNA concentrations determined by UV spectrometry (Chomczynski and Sacchi 2006). Equivalent mass of RNA was used for cDNA synthesis, conducted with Moloney murine leukemia virus reverse transcriptase (Promega) and random hexamers (Roche Applied Science) or iScript cDNA synthesis kit (BioRad). cDNA was diluted five-fold and quantitative PCR (RT-qPCR) was performed using iQ SYBR Green Supermix (Bio-Rad) per manufacturer’s protocol on a 7900HT Sequence Detection System (Applied Biosystems) or with Luna Universal qPCR Master Mix (New England Biolabs) on a CFX Connect Real-Time PCR Detection System (BioRad). RT-qPCR data from polysome fractions was reported as fraction of total mRNA for the indicated gene across the entire gradient (2^-C_t_ for a given fraction divided by the sum of 2^-C_t_ for all fractions). RT-qPCR data from total cell RNA samples was presented as indicated gene mRNA level relative to GAPDH mRNA level (2^-ΔC_t_). Primers used include *GRP94*, CTGGAAATGAGGAACTAACAGTCA (forward) and TCTTCTCTGGTCATTCCTACACC (reverse); *GRP78*, CAACCAACTGTTACAATCAAGGTC (forward) and CAAAGGTGACTTCAATCTGTGG (reverse); *B2M*, TTCTGGCCTGGAGGCTATC (forward) and TCAGGAAATTTGACTTTCCATTC (reverse); *GAPDH*, TCATCAGCAATGCCTCCTGC (forward) and GATGGCATGGACTGTGGTCA (reverse); *XBP1*, CAGCACTCAGACTACGTGCA (forward) and ATCCATGGGGAGATGTTCTGG (reverse); *XBP1*-spliced, CTGAGTCCGAATCAGGTGCAG (forward) and ATCCATGGGGAGATGTTCTGG (reverse).

### Cell fractionation

Sequential detergent cell fractionation was performed as described in (Jagannathan et al. 2011). Briefly, cell cultures were washed with ice-cold PBS, incubated on ice for 20 minutes, and permeabilized with a cytosol extraction buffer containing 110 mM KCl, 25 mM K-HEPES, pH 7.2, 2.5 mM MgOAc_2_, 1 mM EGTA, 1 mM DTT, 50 μg/mL cycloheximide, 1X protease inhibitor cocktail, 40 U/mL RNaseOUT, and 0.015% digitonin for 10 min on ice. Digitonin-treated cells were then washed in an identical buffer with the digitonin concentration reduced to 0.004%. The ER membrane was subsequently solubilized by addition of a detergent extraction buffer containing 200 mM KCl, 25 mM K-HEPES, pH 7.2, 10 mM MgOAc_2_, 2 mM DTT, 50 μg/mL cycloheximide, 1X protease inhibitor cocktail, 40 U/mL RNaseOUT, and 2% DDM on ice for 10 minutes. Samples were then fixed and processed for smFISH and/or immunofluorescence as described below.

### Single molecule RNA fluorescence *in situ* hybridization and immunofluorescence

Single molecule mRNA fluorescence *in situ* hybridization (smFISH) was performed with Stellaris FISH probes (LCG Biosearch Technologies) as per the manufacturer’s recommendations. All reagents were prepared in DEPC-treated deionized water. Cell cultures were fixed in 3.7% paraformaldehyde in PBS for 15 min at room temperature, washed twice in PBS, and permeabilized in ice-cold 70% ethanol at 4°C. Following fixation, cells were washed in 2X saline-sodium citrate (SSC) buffer containing 10% deionized formamide (VWR) and hybridized with fluorescently labeled oligonucleotide probe mixture for 4-16 hours at 37°C in hybridization buffer (2X SSC, 10% deionized formamide, 10% wt/vol dextran sulfate). At the end of the probe hybridization period, cells were washed twice for 30 min in 2X SSC plus 10% deionized formamide at 37°C with DAPI included in the second wash. Following equilibration in 2X SSC, processed coverslips were mounted onto slides using VectaShield Antifade Mounting Medium (Vector Laboratories).

Combined immunofluorescence and smFISH co-staining studies were performed using the protocol recommended by LCG Biosearch Technologies for sequential Stellaris FISH and immunofluorescence for adherent cell cultures. Briefly, cell cultures were fixed as above and permeabilized in 0.1% Triton X-100/PBS for 5 minutes on ice, followed by two sequential washes in ice-cold PBS. For cells that had undergone sequential detergent fractionation, the Triton X-100 permeabilization step was omitted. Detergent-permeabilized cells were blocked in 1% RNAse-free UltraPure BSA (Ambion) for 1 hour at room temperature, stained for 1 hr in primary antibody diluted in 1% BSA/PBS, washed with PBS, stained for 30 min in secondary antibody diluted in 1% BSA/PBS, washed with PBS, and post-fixed in 100% cold methanol for 5 min on ice. Cells were then washed with 2X SSC plus 10% deioninzed formamide, hybridized with fluorescent smFISH probes for 4-16 hours at 37°C, washed, equilibrated, and mounted as described above. Antibodies used include HuR (a gift from Dr. Jack Keene, Duke University School of Medicine), G3BP1 (Santa Cruz, Cat. No. sc-365338), PABP (a gift from Dr. Jack Keene, Duke University School of Medicine), TRAPα (Migliaccio et al. 1992), β-tubulin (Developmental Studies Hybridoma Bank, Cat. No. E7), Alexa Fluor® 488 goat anti-mouse (Thermo Fisher, Cat. No. A11001), Alexa Fluor® 555 goat anti-mouse (Thermo Fisher, Cat. No. A21422), Alexa Fluor® 488 goat anti-rabbit (Thermo Fisher, Cat. No. A11008), Alexa Fluor® 555 goat anti-rabbit (Thermo Fisher, Cat. No. A21428), and Alexa Fluor® 647 goat anti-mouse (Thermo Fisher, Cat. No. A 21235).

### Fluorescence imaging and image processing

All imaging experiments were performed on a DeltaVision Elite deconvolution microscope (Applied Precision) equipped with 100X NA 1.4 oil immersion objective (UPlanSApo 100XO; Olympus) and a high-resolution CCD camera (CoolSNAP HQ2; Photometrics) and conducted at the Duke Light Microscopy Core Facility. Images were acquired as Z-stacks (0.2 μm intervals) at identical exposure conditions across samples for a given protein or smFISH probe. The data were deconvolved using the SoftWoRx program (v6.1 with system-level queuing) (Applied Precision) and processed on ImageJ/FIJI software (Schindelin et al. 2012) to merge channels and pseudo-color images or Imaris 9.6 software for 3D renderings. Only linear changes were made to the brightness/contrast values of the images as required for visualization of patterns and distributions. For smFISH data, brightness/contrast settings were adjusted to ensure optimal visualization of the RNA molecules, while not altering the number of foci in a given sample.

## Acknowledgments

The authors are grateful to Molly Hannigan and JohnCarlo Kristofich for critical comments and feedback on the manuscript and to Alyson Hoffman for scientific contributions to the early phases of the project. This work was supported by grants from the NIH (GM101533, GM118630, CVN) and by shared instrumentation grant NIH 1S10RR027867 (DeltaVisionElite).

## Author contributions

Jessica R. Child; conceptualization, data curation, formal analysis, investigation, methodology, visualization, writing – original draft, review, and editing. Qiang Chen; investigation, data curation, visualization. David W. Reid; data curation, formal analysis, software, visualization, writing – review and editing. Sujatha Jagannathan; conceptualization, formal analysis, investigation, methodology, writing – review and editing. Christopher V. Nicchitta; conceptualization, formal analysis, funding acquisition, methodology, supervision, writing – original draft, review and editing.

## Figure Legends

**Movie 1**. 3D rendering of intact cell with 180° view of SGs corresponding to **Fig. 2D** (unfractionated) depicting GRP94 smFISH (magenta) and immunostaining for the ER membrane protein TRAPα (yellow) and stress granule protein marker G3BP1 (cyan). DAPI staining (blue) identifies the nucleus. TRAPα and G3BP1 are represented as transparent objects to enable visualization of granule association with the ER membrane and co-localization of GRP94 mRNA with G3BP1.

**Movie 2**. 3D rendering of digitonin-permeabilized cell with 180° view of SG corresponding to **Fig. 2D** (digitonin) depicting GRP94 smFISH (magenta) and immunostaining for the ER membrane protein TRAPα (yellow) and stress granule protein marker G3BP1 (cyan). DAPI staining (blue) identifies the nucleus. TRAPα and G3BP1 are represented as transparent objects to enable visualization of granule association with the ER membrane and co-localization of GRP94 mRNA with G3BP1.

